# RADF: Reference-Anchored Dynamic Flow for Spatial Perturbation Profile Completion

**DOI:** 10.64898/2026.08.20.745474

**Authors:** Hongmin Cai, Wanghou, Jiazhou Chen, Zhengfa Xue, Tinghe Zhang, Xiaoqi Sheng

## Abstract

Spatial perturbation profiling is becoming an important tool in functional genomics because it reveals how genetic interventions reshape transcription within intact tissue contexts. However, destructive readout and limited screening capacity leave many perturbation-by-location response profiles unmeasured, motivating the task of spatial perturbation profile completion. The task is to infer the held-out response population at query locations from reported profiles of the same perturbation. Existing methods either generate responses de novo or reuse these profiles without spatial adaptation. These strategies make it difficult to preserve empirical population structure while modeling location-specific variation. Our key insight is that the reported population already defines an empirical response distribution for the target perturbation. To exploit this empirical support, we propose Reference-Anchored Dynamic Flow (RADF), which employs a Sinkhorn-balanced decoder to construct a population-valued anchor in which every reference profile has equal total contribution. Additionally, a bounded dynamic relational flow is used to recompute spatial relations from the evolving expression state and query geometry. Across diverse spatial contexts, RADF reduces macro E-distance by 70.6% compared with an existing state-of-the-art spatial method, highlighting the advantage of combining a reference-supported population anchor with bounded, location-dependent refinement. Code will be made publicly available upon acceptance.

## Introduction

Spatial perturbation profiling combines genetic intervention with spatially resolved molecular measurements, enabling researchers to observe how perturbations reshape transcription within spatially organized tissues rather than only in isolated cells. Spatial perturbation profiling combines genetic intervention with spatially resolved molecular measurements, revealing how perturbations reshape transcription within intact tissues. Recent spatial CRISPR platforms demonstrate that the same intervention can produce distinct response distributions across tissue locations (Dhainaut et al. 2022; Binan et al. 2025; Zhang et al. 2026; Shen et al. 2026; Baysoy et al. 2026; Hu et al. 2025). Because the underlying assays use destructive readout and cannot exhaustively cover every perturbation-by-location population, computational completion of missing responses is essential (Chi et al. 2026; Hu et al. 2025).

Existing perturbation-prediction methods leave this setting only partially addressed. Dissociated-cell models include latent-shift and compositional approaches such as scGen and CPA, graph-based predictors such as GEARS, and distributional mappings such as CellOT and recent flow-based methods (Lotfollahi, Wolf, and Theis 2019; Lotfollahi et al. 2023; Roohani, Huang, and Leskovec 2024; Bunne et al. 2023; Chi et al. 2026). Although these methods differ substantially in architecture, they are designed primarily for dissociated populations and do not explicitly model relations among tissue locations. In our task-adapted implementations, they process spots independently, so their predictions do not use the joint arrangement of query coordinates. Spatial perturbation models incorporate tissue organization more directly: SpatialProp uses graph neural networks to model tissue-level responses, whereas CONCERT employs a niche-aware generative architecture (Sun et al. 2025; Lin et al. 2025). To our knowledge, neither model explicitly uses a reported same-perturbation population as an inference-time population anchor or provides a structural guarantee of balanced reference contribution and bounded deviation from that observed response population (Sun et al. 2025; Lin et al. 2025).

Reference-based copying offers the opposite trade-off. Empirical-reference and nearest-neighbor baselines retain direct support from observed responses, but they are limited to resampling or copying existing profiles and cannot learn controlled refinements based on the joint geometry of the query locations. The resulting gap lies between fully learned response generation and rigid reference reuse. The intermediate problem of observed-perturbation spatial relocation is therefore formulated as follows: given a reported same-perturbation reference population and a set of query coordinates, predict the held-out response distribution while preserving empirical population structure, allowing location-dependent adaptation, and preventing unsupported departures from the observed response support.

We propose **Reference-Anchored Dynamic Flow (RADF)**, a reference-grounded framework for observed-perturbation spatial relocation. RADF separates population support from spatial adaptation. A Sinkhorn-balanced decoder (Cuturi 2013) first allocates reported reference mass across the requested output slots to construct a population-valued anchor. A dynamic spatial relation operator then combines the evolving expression state with query coordinates, and its conditional velocity field updates the anchor only inside a fixed element-wise tube. Validation-only calibration selects the residual strength and checkpoint, returning the exact anchor whenever additional adaptation is unnecessary. Training uses optimal-transport conditional flow matching (Lipman et al. 2023; Liu, Gong, and Liu 2023) and reference E-distance without accessing held-out evaluation expression.

Across matched Perturb-map tasks, RADF consistently outperformed CONCERT and adapted single-cell baselines under both random and spatial-block partitions, and achieved the lowest E-distance among learned methods on sealed same-slide SPAC-seq. The main contributions are:

- **Access-controlled task formulation**. We formulate observed-perturbation spatial relocation with an explicit target-reference access contract and strict separation of fitting, validation, and held-out evaluation expression.
- **Reference-supported population anchoring**. We introduce **RADF**, whose Sinkhorn-balanced reference decoder preserves reference mass while constructing a population-valued response anchor.
- **Bounded geometry-conditioned refinement**. We develop a bounded dynamic spatial residual flow that couples evolving expression states with query geometry and provides an exact anchor fallback.
- **Matched and transfer evaluation**. Extensive matched, cross-slide, and sealed evaluations show that RADF surpasses spatial and task-adapted single-cell predictors in settings most aligned with spatial response completion.

### Related Works and Preliminaries

RADF lies between perturbation prediction and spatial distribution completion. We first locate it among existing models, then define the access contract, population metric, and anchored flow used below.

### Existing Perturbation Prediction Models

Perturbation predictors take several routes: scGen and CPA learn latent or compositional effects, GEARS uses graph-based regression, and CellOT or flow-based models map entire distributions (Lotfollahi, Wolf, and Theis 2019; Lotfollahi et al. 2023; Roohani, Huang, and Leskovec 2024; Bunne et al. 2023; Chi et al. 2026). Their usual target is a dissociated cell population. In our Visium adaptations, they process spots independently; rearranging the same query spots in tissue leaves their predictions unchanged. They also lack RADF’s explicit contract in which a reported population from the same perturbation supplies the response support.

SpatialProp and CONCERT do model tissue-level receiver effects or niche-dependent perturbation representations (Sun et al. 2025; Lin et al. 2025). Their end-to-end construction still has to recover response support and context variation together. RADF uses the available information differently: the target-reference population supplies empirical support, and learning is reserved for a bounded change across query contexts.

### Problem Definition

Let *p* denote an observed perturbation and *G* the number of aligned genes. Its measured target pool contains perturbed expression vectors and spatial coordinates. Before model fitting, this pool is partitioned using row identities or coordinates only into a reported reference set, a validation set, and a held-out evaluation set:

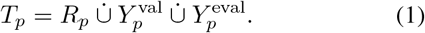

Here 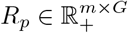, and the three subsets in Equation (1) are mutually disjoint. For *n* requested outputs, the observable spatial input is *S* = [*s*_1_, …, *s*_*n*_]^⊤^ ∈ ℝ^*n×*2^. RADF predicts a nonnegative expression population of the same output size:

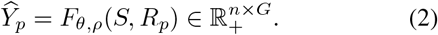

The perturbation label indexes the task and its reported reference population, and RADF is fitted separately for each observed intervention. The predictor completes an observed response distribution rather than reconstructing paired cells or predicting an intervention for which no same-perturbation reference is available.

Fitting and candidate generation may read only (*S, R*_*p*_). Candidate parameters and predictions are frozen before 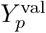 is opened. Validation E-distance selects the residual radius and checkpoint, including an anchor fallback, after which the selection is frozen. 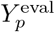 is read only once for final scoring and cannot affect fitting or selection.

### E-Distance and Anchored Flow

Because predicted and observed cells are unpaired and their population sizes may differ, the primary objective is empirical E-distance (Székely and Rizzo 2013). Its empirical V-statistic is stated in the supplementary material and uses Euclidean distance in aligned gene space. Evaluation clamps only negative numerical round-off to zero; lower values are better. Metrics are computed per task/split unit and macro-averaged so that larger target pools do not dominate. Because E-distance compares unordered populations, it is invariant to output-row permutations and cannot by itself establish coordinate sensitivity.

RADF begins from an anchor population 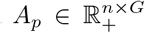 produced from *R*_*p*_. Both the learned anchor weights and the fixed training coupling use balanced matrices (Cuturi 2013) in

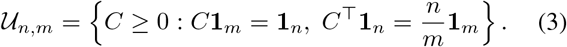

Equivalently, *C/n* is a unit-mass balanced coupling with uniform marginals 1*/n* and 1*/m*; meanwhile, non-negativity and *C***1**_*m*_ = **1**_*n*_ make each row of *CR*_*p*_ a convex combination of reference profiles, while the column constraint assigns every reference equal total contribution *n/m*. For a fixed entropic optimal-transport coupling Γ_*p*_ ∈ *U*_*n,m*_ between *A*_*p*_ and *R*_*p*_, the coupled endpoint is *B*_*p*_ = Γ_*p*_*R*_*p*_. Conditional flow matching uses the straight path and its constant target velocity (Lipman et al. 2023; Liu, Gong, and Liu 2023):

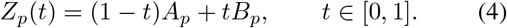

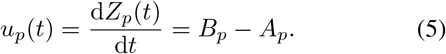

The learned field *v*_*θ*_(*Z*_*p*_(*t*), *t*; *S, R*_*p*_) approximates this velocity. At inference, projected Euler updates keep every state inside the elementwise anchor tube

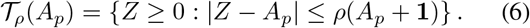

Consequently, zero velocity returns *A*_*p*_ exactly, while every nonzero contextual refinement is explicitly bounded.

## Methods

### Overview

RADF has a deliberately asymmetric pipeline. The reported set *R*_*p*_ carries response support; the learned flow handles only its spatial adjustment. A Sinkhorn-balanced allocation first produces *n* anchor profiles *A*_*p*_, with fixed total contribution from every reference. A conditional velocity network then rebuilds relations from the evolving expression and query coordinates *S*, followed by projected Euler updates inside an anchor tube. Finally, validation Edistance chooses the radius and checkpoint, or keeps *A*_*p*_ when none of the learned updates helps. The prediction map is 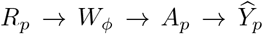, with 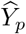 conditioned on (*S, R*_*p*_, *t*). All transformations remain in expression space. Coordinates organize relations among output slots but are not themselves transported. Figure 1 follows the same division through training, validation-only selection, and frozen inference.

**Figure 1:**
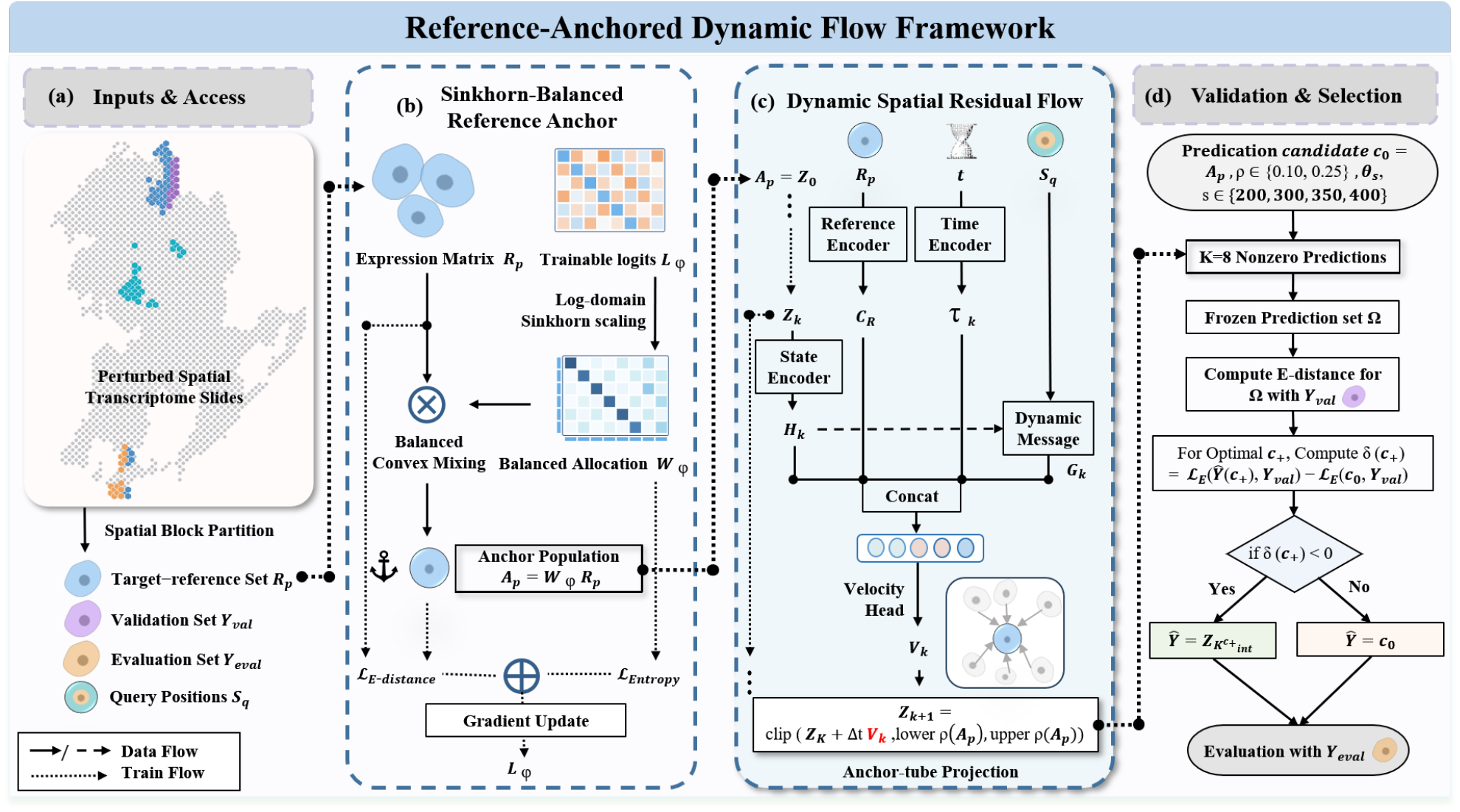
Overview of RADF. (a) Spatial-block partitioning separates the target-reference, validation, and held-out evaluation sets, with query positions provided as spatial context. (b) Sinkhorn-balanced convex allocation transforms reference profiles into the anchor population *A*_*p*_. (c) Starting from *A*_*p*_, dynamic spatial messages, reference context, and time determine bounded velocity updates within the anchor tube. (d) Validation E-distance selects among frozen radius–checkpoint candidates, retaining a nonzero flow only when it improves upon the anchor before final evaluation.

### Sinkhorn-Balanced Reference Anchor

Let 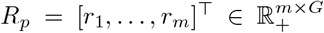, and let *n* be the requested number of output samples. The decoder maintains unconstrained trainable logits *L*_*ϕ*_ ∈ ℝ^*n×m*^. Log-domain Sinkhorn scaling converts exp(*L*_*ϕ*_) to a plan with uniform row and column marginals (Cuturi 2013). Alternating row and column log-normalizations remain differentiable with respect to *L*_*ϕ*_; after scaling the unit-mass plan by *n*, the resulting allocation matrix satisfies *W*_*ϕ*_ ∈ *U*_*n,m*_, with the balance constraints already defined in Equation (3). The exact log-domain recurrences are recorded in the separate supplementary document.

Each row is a probability vector over references, and each reference contributes total mass *n/m* across the output population. Their nonnegative convex allocation is

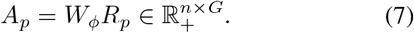

Every anchor row lies in the convex hull of the reported expressions. Row *i* is an output slot, not a biological cell paired to *r*_*i*_. Associating that slot with query coordinate *s*_*i*_ lets the relational flow adjust the population by context without inventing its support.

The allocation logits are learned from reported reference expression alone by minimizing

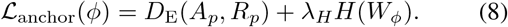

We use stabilized negative entropy,

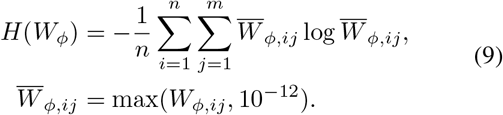

Here *λ*_*H*_ = 0.05. Because *H* enters with a positive coefficient and the objective is minimized, it discourages overly diffuse row allocations. Without this term, convex mixtures can smooth distinct references into similar outputs while still matching their center. The column constraint prevents sharper rows from dropping any reference cell at the population level. This is an allocation regularizer, not a biological similarity prior. The implementation uses 160 Sinkhorn iterations at each decoder update and fits the logits for 400 Adam steps with learning rate 0.05.

Balance provides the exact population-mean identity 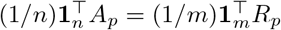. Thus the anchor preserves the reference mean and uniform reference contribution by construction, including when *n* differs from *m*. Its *n* rows retain a population-valued representation rather than collapsing to one centroid. This identity preserves allocation over reference samples, not biological RNA mass, total expression, or gene-wise library size; throughout, balanced therefore refers only to the empirical sample measure represented by *R*_*p*_. The flow then provides bounded query-context refinement.

### Bounded Dynamic Relational Flow

Let 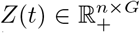 be the evolving population, initialized by *Z*(0) = *A*_*p*_. RADF parameterizes a conditional velocity field

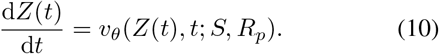

The field recomputes its relation matrix from the current state at every integration step. For row *z*_*i*_(*t*), the state encoder is

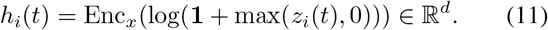

Query coordinates are centered and scaled per axis, with the scale lower-bounded by 10^−3^:

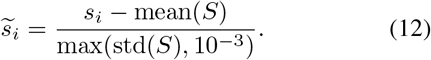

With learned linear maps *Q*_*h*_ and *K*_*h*_, let *q*_*i*_(*t*) = *Q*_*h*_*h*_*i*_(*t*) and *k*_*j*_(*t*) = *K*_*h*_*h*_*j*_(*t*). The off-diagonal relation scores are

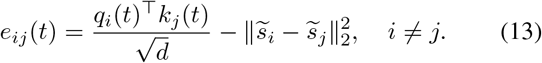

For *n >* 1, self-scores are masked by setting *e*_*ii*_(*t*) = −10^9^; the normalized weights are

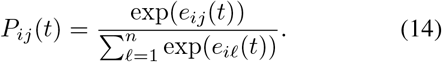

The spatial penalty has fixed coefficient one after coordinate standardization. Because *Q*_*h*_ and *K*_*h*_ are distinct and normalization is row-wise, *P* (*t*) is generally directed, dense, and asymmetric. Consequently, *P* (*t*)**1**_*n*_ = **1**_*n*_, and the message *g*_*i*_(*t*) = ∑_*j*_ *P*_*ij*_(*t*)*h*_*j*_(*t*) is a row-stochastic aggregation. For a single output, the unique relation weight is set to one. Since *h*_*i*_(*t*) changes with *Z*(*t*), *P* (*t*) is state-dependent even though the coordinates remain fixed.

The reported reference set enters through a global distributional summary. Let *µ*_*R*_ and *σ*_*R*_ be its gene-wise mean and standard deviation. We encode them as *c*_*R*_ = Enc_*R*_(log(**1** + max(concat(*µ*_*R*_, *σ*_*R*_), 0))), while *τ*_*i*_(*t*) = Enc_*t*_(*t*) encodes time. The same *c*_*R*_ is broadcast to all output slots, providing a shared summary of the observed response distribution. Slot-specific adaptation is then determined by the evolving state, its spatial message, and continuous time.

The velocity head combines the local state, its dynamic relational message, the shared reference summary, and time:

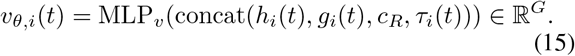

The state encoder maps *G* genes to *d* features, the reference encoder maps 2*G* moments to *d*, and the time encoder maps one scalar to *d*. The velocity head maps the concatenated 4*d* features through a 2*d* GELU hidden layer to *G* outputs. All three encoders use GELU activations; the state and reference encoders additionally use layer normalization. The benchmark uses *d* = 32. The final linear layer of MLP_*v*_ is initialized to zero, so the untrained residual path initially returns the anchor.

Rather than solving an unrestricted neural ODE (Chen et al. 2018), RADF uses *K*_int_ projected Euler steps. The fixed elementwise limits in Equation (6) are

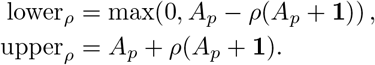

The additive one permits a bounded update even for genes whose anchor value is near zero, while the multiplicative part scales the allowed deviation with expression magnitude. These bounds are fixed from *A*_*p*_ and are not recomputed from later states.

With *Z*_0_ = *A*_*p*_, Δ*t* = 1*/K*_int_, and midpoint time 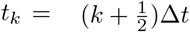, inference applies

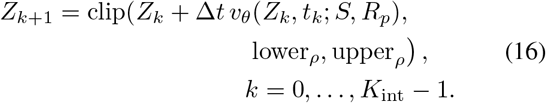

The final prediction is 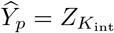. Clipping is elementwise and is performed after every step; *K*_int_ = 8 in the benchmark. Hence every finite prediction is nonnegative and remains inside the chosen anchor tube. Zero velocity leaves *Z*_*k*_ = *A*_*p*_, giving an exact structural fallback; nonzero updates are coordinate-conditioned relational refinements in expression space.

### Training and Conservative Selection

RADF trains the residual field against a fixed balanced endpoint derived from *A*_*p*_ and *R*_*p*_. We standardize their pooled expression gene-wise using

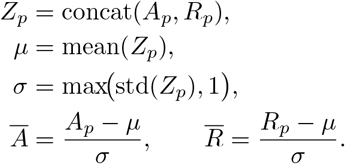

The coupling cost is 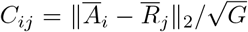. Log-domain Sinkhorn scaling of −*C/ϵ*, with *ϵ* = 0.5 and 120 scaling iterations, produces a fixed Γ_*p*_ ∈ *U*_*n,m*_, and *B*_*p*_ = Γ_*p*_*R*_*p*_. This endpoint supervises reference-supported distributional refinement. Because both *A*_*p*_ and *B*_*p*_ are derived from *R*_*p*_, the CFM target is not location-supervised; coordinates influence the field only through Equations (13)–(14).

For independently sampled *t*_*i*_ ∼ Uniform(0, 1), define *z*_*i*_(*t*_*i*_) = (1 − *t*_*i*_)*a*_*i*_ + *t*_*i*_*b*_*i*_ and *u*_*i*_ = *b*_*i*_ − *a*_*i*_. The detached normalization scales are

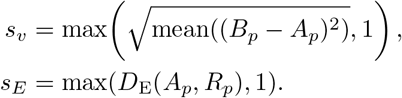

Training minimizes the normalized conditional flow-matching loss and the E-distance of the projected prediction to the reported references (Lipman et al. 2023; Liu, Gong, and Liu 2023):

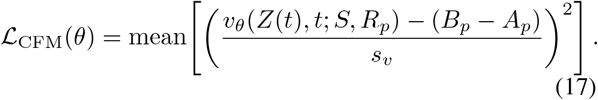

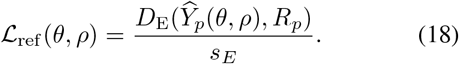

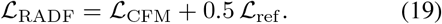

The fixed coupling, anchor, and scales are detached from gradient updates. Models are optimized for 400 Adam steps with learning rate 10^−3^, with gradient norm clipped to five. The differentiable algebraic E-distance is used during optimization without the evaluation-only nonnegative clamp. Separate models with the same deterministic initialization are trained for *ρ* ∈ {0.10, 0.25} because projection changes ℒ_ref_ ; predictions and states are saved at checkpoints 200, 300, 350, 400. Together with the anchor candidate *c*_0_, this yields a frozen candidate set before validation expression is read.

For validation set 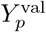, define each candidate’s improvement as

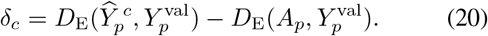

The best nonzero candidate is

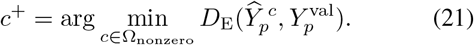

Conservative selection chooses

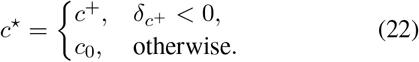

Ties among nonzero candidates prefer the smaller radius and then the earlier checkpoint. This strict comparison implements conservative selection: a residual is retained only when validation supports improvement over zero residual. The selected prediction is frozen before held-out evaluation expression is opened. PCC and MAE are reported descriptively but do not enter training, selection, or fallback.

## Experiment

Evaluation covers matched Perturb-map tasks, adapted single-cell predictors, cross-slide transfer, and sealed SPAC-seq. E-distance is the primary endpoint. Mean PCC correlates predicted and observed gene-wise population means, and mean MAE measures their absolute difference. For spatial diagnostics, pooled query and truth coordinates are divided at the median projection onto their first spatial principal axis. SS-ED averages the two within-stratum E-distances with truth-count weights; spatial contrast error compares the predicted and observed separation between strata. Each stratum must contain at least four samples.

### Experimental Setup

The matched Perturb-map benchmark (Dhainaut et al. 2022) contained five slide-perturbation tasks: Tgfbr2 on GSM5808054, GSM5808055, and GSM5808057, and Ifngr2 on GSM5808055 and GSM5808056. Each task used the 31 nearest KP/tumor query spots and 2,000 ordered genes in frozen raw-count space. Target rows were split into 60% references, 20% validation, and the remainder evaluation. Random splits used seed 20260731; spatial-block splits followed coordinate order, producing ten units. Official CONCERT used its fixed multi-kernel implementation after heldout rows were removed from its fit file (Lin et al. 2025). CellOT, scGen, and biolord used fixed configurations with the same source, reference, genes, and held-out-expression access (Bunne et al. 2023; Lotfollahi, Wolf, and Theis 2019; Piran et al. 2024).

External experiments added five leave-one-slide-out Perturb-map units on 1,053 shared genes, each with 32 references from other slides. SPAC-seq (Zhang et al. 2026) used the released expression matrix on 1,718 genes intersecting the frozen panel for sgZc3h12a, sgZhx2, and sgGata3 across days 4, 7, and 10. The sealed same-slide track contained 18 units with 32 references and 64 disjoint evaluation spots; the cross-replicate track contained nine units with 64 replicate-1 references and 128 replicate-2 evaluation spots. Radius 0.25 and checkpoint 300 were frozen from Perturb-map validation. Absolute E-distances are interpreted only within a dataset and preprocessing space.

Metrics were computed independently for each unit and then macro-averaged. Paired intervals used 10,000 clusterbootstrap draws over tasks, held-out slides, or days. These intervals describe cross-unit consistency; no spot-level significance tests were used.

### RADF Outperforms CONCERT on Matched Tasks

RADF has lower E-distance than CONCERT in every matched unit (Table 1). Overall macro error falls from 138.05 to 40.60, a reduction of 70.60%. The reduction is 80.70% on random splits and 63.20% on spatial blocks; the task-cluster interval for CONCERT minus RADF is [64.39, 132.39]. Population means improve as well: PCC rises from 0.92 to 0.97 and MAE drops from 0.68 to 0.38. The scale and consistency of this gap favor a reference-supported predictor over relearning the full response distribution with a general conditional generator.

**Table 1:** Matched Perturb-map results reported as mean *±* population standard deviation across task/split units (overall *n* = 10; each partition *n* = 5). Arrows indicate the optimization direction. Bold and underlined means denote the best and second-best results, respectively.

| Method | Overall |  |  | Random |  |  | Spatial block |  |  |
| --- | --- | --- | --- | --- | --- | --- | --- | --- | --- |
|  | E-dist ↓ | PCC ↑ | MAE ↓ | E-dist ↓ | PCC ↑ | MAE ↓ | E-dist ↓ | PCC ↑ | MAE ↓ |
| CONCERT | 138.05 ± 55.74 | 0.92 ± 0.04 | 0.68 ± 0.26 | 117.15 ± 28.02 | 0.94 ± 0.02 | 0.59 ± 0.19 | 158.95 ± 67.50 | 0.90 ± 0.05 | 0.77 ± 0.28 |
| scGen | <u>62.98</u> ± 34.29 | <u>0.96</u> ± 0.03 | <u>0.43</u> ± 0.15 | <u>43.79</u> ± 14.79 | <u>0.98</u> ± 0.01 | <u>0.33</u> ± 0.07 | <u>82.17</u> ± 37.36 | <u>0.94</u> ± 0.03 | <u>0.53</u> ± 0.14 |
| biolord | 102.02 ± 41.26 | 0.96 ± 0.03 | 0.46 ± 0.15 | 86.65 ± 21.72 | 0.97 ± 0.02 | 0.39 ± 0.09 | 117.40 ± 49.59 | 0.94 ± 0.04 | 0.54 ± 0.17 |
| CellOT | 461.26 ± 250.84 | 0.76 ± 0.08 | 1.97 ± 0.50 | 447.28 ± 307.12 | 0.78 ± 0.09 | 1.93 ± 0.58 | 475.25 ± 176.43 | 0.74 ± 0.07 | 2.00 ± 0.42 |
| **RADF (Ours)** | **40.60** ± 23.92 | **0.97** ± 0.02 | **0.38** ± 0.12 | **22.64** ± 11.02 | **0.99** ± 0.01 | **0.29** ± 0.05 | **58.55** ± 19.46 | **0.96** ± 0.01 | **0.48** ± 0.10 |

The same separation is visible in Figure 2. Overall Edistance is 1.55 times RADF’s error for the strongest adapted single-cell baseline, 3.40 times for CONCERT, and 11.36 times for CellOT.

**Figure 2:**
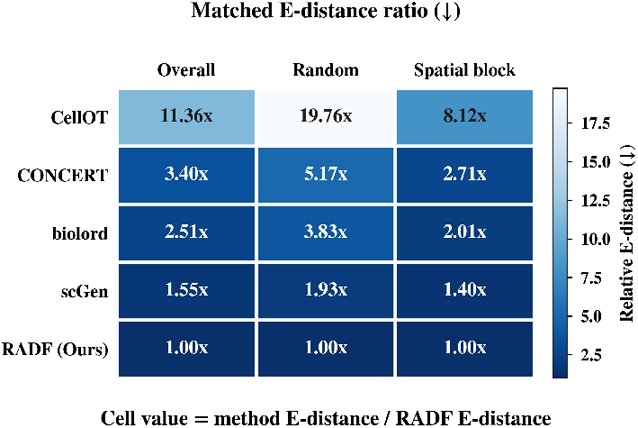
Matched Perturb-map E-distance ratios. Each cell is a method’s E-distance divided by RADF E-distance for the same partition; lower is better and values above one indicate higher error.

### Why Single-Cell Predictors Fall Short

Spatial relocation lies outside the native design of singlecell perturbation predictors. scGen applies a shared latent displacement, biolord models condition attributes without tissue relations, and CellOT maps dissociated populations without query coordinates. Their Visium adaptation treats spots as independent pseudo-cells. Two query sets with the same spot profiles but different tissue arrangements therefore receive the same prediction. RADF instead retains the observed response population and rebuilds relations over the supplied query geometry.

That difference in task design is reflected in the matched results. RADF records 40.60 overall E-distance, compared with 62.98 for scGen, 102.02 for biolord, and 461.26 for CellOT, and wins all ten units against each method. Equal expression and reference access is therefore not enough when the predictor has no mechanism to relocate a population across tissue contexts.

The advantage transfers beyond the matched split. RADF cuts leave-one-slide-out CONCERT error from 155.30 to 71.20. On sealed same-slide SPAC-seq, it is the strongest learned method at 0.47 E-distance, against 3.69 for CONCERT and 1.15 for scGen. Under cross-replicate variation, RADF remains ahead of CONCERT and has the lowest spatial contrast error.

### What the Flow Contributes

The anchor-versus-RADF average measures the realized marginal value of validation-gated refinement above an already strong anchor; it does not measure the representational capacity of the flow. Figure 3 separates these questions. In a matched-architecture development audit over four earlier units, a relational flow reached 35.92 E-distance versus 35.67 for the reference decoder, a gap of 0.70%. On two untouched prospective units, an anchored relational flow reduced E-distance by 1.24% on a spatial split and 2.91% on a random split. Because the three rows use different protocols, they are shown separately and are not pooled with the paper benchmark.

**Figure 3:**
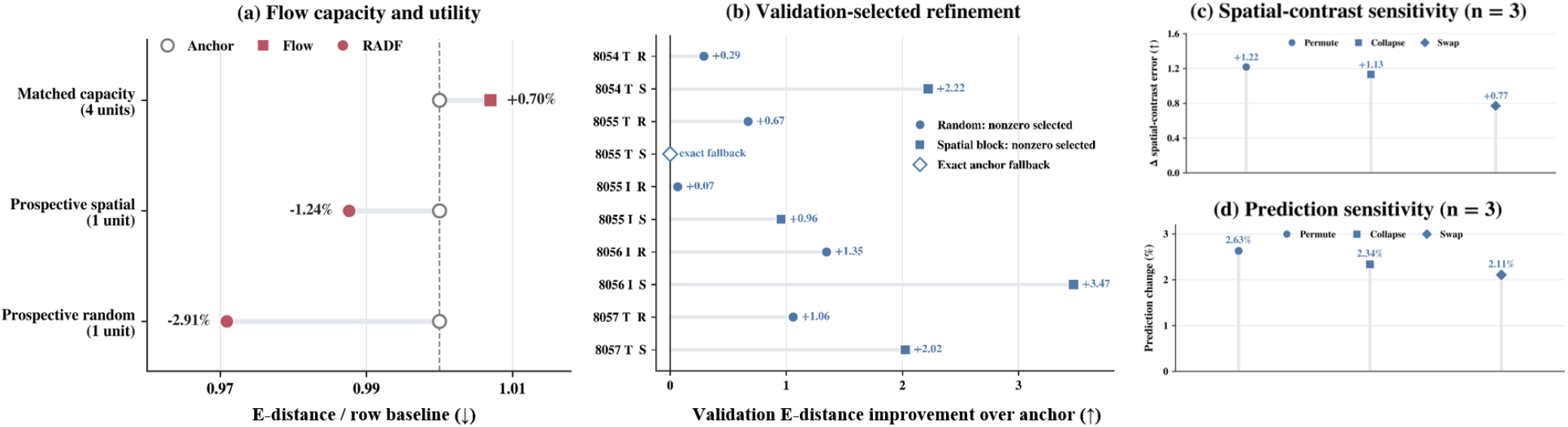
Flow capacity, selective utility, and spatial sensitivity. (a) Row-normalized E-distance: capacity compares flow with a matched reference decoder across four development units; prospective rows compare RADF with its anchor on one spatial and one random unit. Protocols differ and are not pooled. (b) Per-unit gain over the anchor after validation-only selection; positive values favor RADF. Filled marks indicate nonzero flow; the open square marks exact fallback. T/I denote Tgfbr2/Ifngr2 and R/S random/spatial-block splits. (c) Change in spatial-contrast error under coordinate interventions. (d) Relative prediction change under the same interventions. Panels (c)–(d) use three supported units; supplementary material reports SS-ED and unit-level results.

On the ten matched units, validation selected nonzero flow in nine and returned the exact anchor in one. Selected residuals changed 2.2–7.9% of the anchor magnitude. RADF improved over the learned anchor by 0.24% overall, decomposing into +2.31% on random splits and -0.58% on spatial blocks; per-unit changes ranged from +5.36% to -1.47%. The modest macro gain therefore reflects limited residual headroom after anchoring and task-dependent transfer across heterogeneous spatial blocks. The anchor tube and exact fall-back keep this refinement conservative.

E-distance alone cannot reveal where an output belongs because row permutation leaves it unchanged. Across the three supported units, permuting, collapsing, or swapping coordinates changed predictions by 2.11–2.63% and increased spatial-contrast error by 0.77–1.22. The fitted flow therefore uses spatial inputs rather than acting as a coordinate-invariant decoder. Complementary SS-ED outcomes and all unit-level selection results are reported in the separate supplementary document.

A separate truth-assisted diagnostic found context-specific improvements of 2.25% and 2.24% over its matched global Oracle within a different frozen rank-4, 128-gene family. These are achieved representation-level increments, not a global ceiling or an estimator-to-Oracle gap; full protocol details are provided in the separate supplementary document.

### Transfer Across Spatial Contexts

The advantage over CONCERT remains as evaluation contexts become more separated: RADF reduces E-distance by 80.70% on random partitions, 63.20% on held-out spatial blocks, and 54.2% when an entire slide is left out. Reference anchoring therefore transfers beyond interpolation among nearby spots.

Spatially stratified diagnostics sharpen this picture (Figure 4). On supported matched units, RADF lowers SS-ED from 138.40 for CONCERT to 65.91 and contrast error from 83.30 to 56.00. On sealed same-slide SPAC-seq, the corresponding reductions are 3.91 to 0.84 and 0.65 to 0.20. Cross-replicate contrast error is 0.30, below scGen (0.38), empirical reference prediction (0.38), and CONCERT (0.51). These regional endpoints complement the population-level ranking and isolate the spatial structure most relevant to the task.

**Figure 4:**
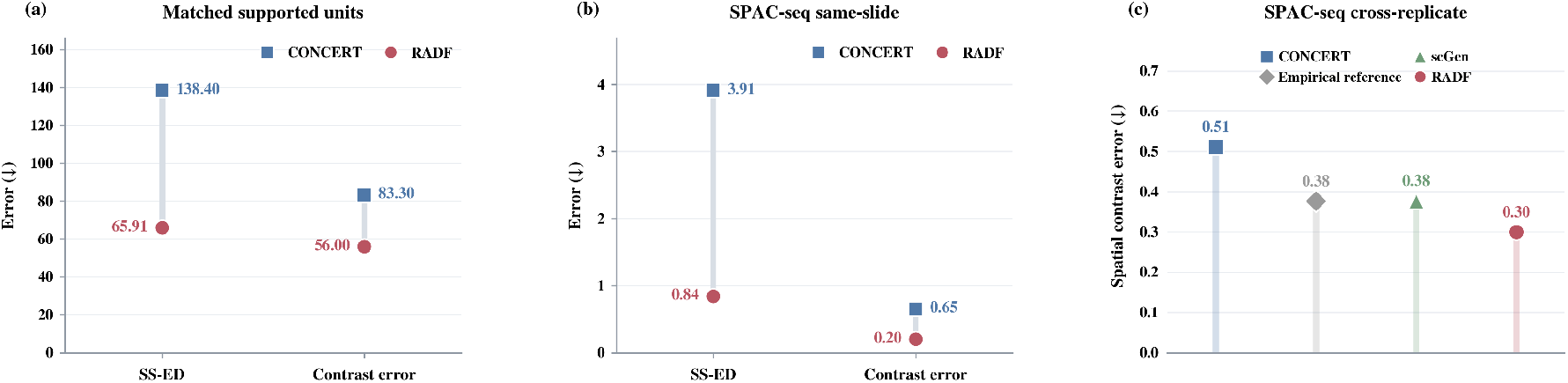
Spatial diagnostics emphasizing RADF’s regional-distribution advantage. (a) Matched supported units. (b) Sealed same-slide SPAC-seq. (c) Cross-replicate spatial contrast error.

RADF also completes cross-slide evaluation in 38.9 seconds, compared with 63.6 for CONCERT, and SPAC-seq in 39.6 seconds, compared with 84.2. Cross-replicate variation is the hardest setting and provides a clear target for slide-aware reference normalization.

## Conclusion

In this study, we proposed RADF to complete unmeasured spatial response profiles for observed perturbations with reported reference populations. It combines a Sinkhorn-balanced reference anchor with a bounded dynamic relational flow. The anchor retains reference-supported population structure, while the flow enables controlled, geometry-conditioned refinement. Across matched and transfer settings, RADF reduces population error and improves regional fidelity. These results establish RADF as a reference-grounded alternative to fully learned spatial response generation.

## Notes

### Competing Interest Statement

The authors have declared no competing interest.

## References

Baysoy, A.; Tian, X.; Renauer, P.; Zhang, F.; Bai, Z.; Shi, H.; Yang, M.; Zhang, D.; Liu, M.; Li, H.; Tao, B.; Enninful, A.; Lu, Y.; Gao, F.; Wang, G.; Zhang, W.; Tran, T.; Patterson, N. H.; Sheng, J.; Bao, S.; Dong, C.; Xin, S.; Chen, B.; Zhong, M.; Rankin, S.; Guy, C.; Wang, Y.; Connelly, J. P.; Pruett-Miller, S. M.; Wang, D.; Xu, M.; Gerstein, M. B.; Chi, H.; Chen, S.; and Fan, R. 2026. Large-scale, spatially resolved panoramic CRISPR screening in native tissue environments using Perturb-DBiT. Nature Biotechnology. Advance online publication. doi: 10.1038/s41587-026-03127-y.

Binan, L.; Jiang, A.; Danquah, S. A.; Valakh, V.; Simonton, B.; Bezney, J.; Manguso, R. T.; Yates, K. B.; Nehme, R.; Cleary, B.; and Farhi, S. L. 2025. Simultaneous CRISPR screening and spatial transcriptomics reveal intracellular, intercellular, and functional transcriptional circuits. Cell, 188(8): 2141–2158.e18.

Bunne, C.; Stark, S. G.; Gut, G.; del Castillo, J. S.; Levesque, M.; Lehmann, K.-V.; Pelkmans, L.; Krause, A.; and Rätsch, G. 2023. Learning single-cell perturbation responses using neural optimal transport. Nature Methods, 20(11): 1759–1768.

Chen, R. T. Q.; Rubanova, Y.; Bettencourt, J.; and Duvenaud, D. 2018. Neural Ordinary Differential Equations. In Advances in Neural Information Processing Systems.

Chi, C.; Huang, Y.; Xia, J.; Zheng, J.; Liu, Y.; Zang, Z.; and Li, S. Z. 2026. Departures: Distributional Transport for Single-Cell Perturbation Prediction with Neural Schrödinger Bridges. Proceedings of the AAAI Conference on Artificial Intelligence, 40(25): 20544–20552.

Cuturi, M. 2013. Sinkhorn Distances: Lightspeed Computation of Optimal Transport. In Advances in Neural Information Processing Systems.

Dhainaut, M.; Rose, S. A.; Akturk, G.; Wroblewska, A.; Nielsen, S. R.; Park, E. S.; Buckup, M.; Roudko, V.; Pia, L.; Sweeney, R.; Berichel, J. L.; Wilk, C. M.; Bektesevic, A.; Lee, B. H.; Bhardwaj, N.; Rahman, A. H.; Baccarini, A.; Gnjatic, S.; Pe’er, D.; Merad, M.; and Brown, B. D. 2022. Spatial CRISPR genomics identifies regulators of the tumor microenvironment. Cell, 185(7): 1223–1239.e20.

Hu, M.; Cui, Y.; Huang, Q.; Chu, K.; McKinzie, S.; Patrick, M.; Iyengar, S.; Abuduli, M.; Spatz, M.; Joshi, N.; Miller, B.; Vellarikkal, S.; Riordan, T.; Bitton, D.; Lubojacky, J.; Khalil, I.; Piccioni, F.; Rhodes, M.; Tamburino, A.; He, S.; Beechem, J.; and Peterson, V. 2025. SPACE: multimodal spatial CRISPR screening with whole-transcriptome readout at subcellular resolution in 3D models. bioRxiv:2025.09.14.675819.

Lin, X.; Kong, Z.; Ghosh, S.; Kellis, M.; and Zitnik, M. 2025. CONCERT predicts niche-aware perturbation responses in spatial transcriptomics. bioRxiv:2025.11.08.686890.

Lipman, Y.; Chen, R. T. Q.; Ben-Hamu, H.; Nickel, M.; and Le, M. 2023. Flow Matching for Generative Modeling. In International Conference on Learning Representations.

Liu, X.; Gong, C.; and Liu, Q. 2023. Flow Straight and Fast: Learning to Generate and Transfer Data with Rectified Flow. In International Conference on Learning Representations.

Lotfollahi, M.; Susmelj, A. K.; Donno, C. D.; Hetzel, L.; Ji, Y.; Ibarra, I. L.; Srivatsan, S. R.; Naghipourfar, M.; Daza, R. M.; Martin, B.; Shendure, J.; McFaline-Figueroa, J. L.; Boyeau, P.; Wolf, F. A.; Yakubova, N.; Günnemann, S.; Trapnell, C.; Lopez-Paz, D.; and Theis, F. J. 2023. Predicting cellular responses to complex perturbations in high-throughput screens. Molecular Systems Biology, 19(6): MSB202211517.

Lotfollahi, M.; Wolf, F. A.; and Theis, F. J. 2019. scGen predicts single-cell perturbation responses. Nature Methods, 16(8): 715–721.

Piran, Z.; Cohen, N.; Hoshen, Y.; and Nitzan, M. 2024. Disentanglement of single-cell data with biolord. Nature Biotechnology, 42(11): 1678–1683.

Roohani, Y.; Huang, K.; and Leskovec, J. 2024. Predicting transcriptional outcomes of novel multigene perturbations with GEARS. Nature Biotechnology, 42(6): 927–935.

Shen, K.; Seow, W. Y.; Keng, C. T.; Lim, M. G. L.; Lim, D. S.; Guo, K.; Meliani, A.; Hajis, M. I. B.; Wang, B.; Prabhakar, S.; Chen, K. H.; and Chew, W. L. 2026. Spatial perturb-seq: single-cell functional genomics within intact tissue architecture. Nature Communications. Article in press. doi: 10.1038/s41467-026-69677-6.

Sun, E. D.; Buendia, A.; Brunet, A.; and Zou, J. 2025. SpatialProp: tissue perturbation modeling with spatially resolved single-cell transcriptomics. bioRxiv:2025.11.30.691355.

Székely, G. J.; and Rizzo, M. L. 2013. Energy statistics: A class of statistics based on distances. Journal of Statistical Planning and Inference, 143(8): 1249–1272.

Zhang, H.; Zhang, Z.; Wang, P.; Xu, T.; Chen, X.; Zhao, Y.; Lin, S.; Cai, W.; Ren, P.; Luo, C.; Zhang, P.; Wang, Y.; Hou, S.; Zhao, Y.; Zeng, H.; Liu, Z.; Wang, C.; Gao, Z.; Feng, Y.; Pan, D.; and Zeng, Z. 2026. Uncovering spatially resolved functional genomics with CRISPR screen sequencing. Cell, 189(15): 4594–4618.e48.

