## supplementery for "RADF: Reference-Anchored Dynamic Flow for Spatial Perturbation Profile Completion"

### Reference Anchoring Preserves Response Structure

RADF is designed around an asymmetric division of labor: the reported same-perturbation population supplies empirical response support, while the flow is restricted to a bounded spatial refinement. This section isolates the first part of that design. The learned anchor is neither a centroid nor an unconstrained generator. Its feasible allocation set is

$$\mathcal{U}_{n,m} = \left\{ W \geq 0 : W\mathbf{1}_m = \mathbf{1}_n,\ W^\top \mathbf{1}_n = \frac{n}{m}\mathbf{1}_m \right\}. \tag{S1}$$

If $A_p = W_\phi R_p$ and $W_\phi \in \mathcal{U}_{n,m}$, then

$$\frac{1}{n}\mathbf{1}_n^\top A_p = \frac{1}{n}\mathbf{1}_n^\top W_\phi R_p = \frac{1}{m}\mathbf{1}_m^\top R_p. \tag{S2}$$

Thus the anchor preserves the uniform empirical reference mean exactly, while its rows remain convex combinations of observed reference profiles. The constraint controls contribution over reference samples; it does not claim preservation of biological RNA mass or library size.

Table S1 shows why a population-valued anchor is useful. Across the ten matched Perturb-map units, the learned anchor reduces E-distance by 66.0% relative to a repeated uniform reference mean and by 5.2% relative to direct empirical-reference reuse. RADF retains this strong base and reduces the overall mean from 40.693 to 40.595. The small anchor-to-RADF difference should be read as controlled refinement above an already competitive reference representation, not as the total value of the RADF design.

### Bounded Flow Provides Selective Spatial Refinement

The main paper separates three questions that an aggregate anchor comparison cannot answer alone: whether the flow family has sufficient capacity, whether it adds value on untouched units, and whether fitted predictions actually use coordinates. We report these evidence layers separately because they arise from distinct frozen protocols.

#### Matched Capacity and Prospective Utility

The matched-architecture capacity audit compares a relational flow with a reference decoder over four earlier development units. The flow reaches 35.915 E-distance versus 35.666 for the decoder, a ratio of 1.007. This result supports reference-level distributional capacity; it does not claim that the flow is superior to the decoder. Two untouched prospective units then test the full anchored construction. RADF improves over its anchor by 1.24% on the spatial unit and 2.91% on the random unit (Table S2).

#### Validation Retains Useful Flow and Rejects Unsupported Updates

The paper benchmark freezes two nonzero radii and four checkpoints before validation expression is opened. Validation selects a nonzero candidate in nine of ten units and returns the anchor exactly in the remaining unit. The selected evaluation residuals have relative Frobenius magnitude 2.24–7.85%, showing that nonzero selections are bounded corrections rather than wholesale regeneration. Table S3 gives the complete selection receipt. Evaluation favors RADF over the anchor in five units, ties in one, and favors the anchor in four. This 5/1/4 pattern makes the interpretation precise: validation gating yields selective marginal utility, while the exact fallback prevents an unsupported residual from being forced into every task.

#### Coordinate Interventions Confirm Spatial Dependence

E-distance treats predictions as unordered populations and therefore cannot by itself establish coordinate use. We intervene on coordinates while keeping the trained model, reference population, and all other inputs fixed. The prespecified common-stratum rule requires at least four truth and prediction samples in each region; three of ten units satisfy it. On these supported units, permutation, coordinate collapse, and region swap change predictions by 2.11–2.63% and increase spatial-contrast error by 0.77–1.22 (Table S4). The fitted flow is therefore not coordinate invariant.

#### Truth-Assisted Context Headroom

A separate truth-assisted representation diagnostic tests whether a fixed low-rank context component can improve upon a matched global Oracle. The basis, rank, bounds, and optimization budget are frozen before target truth is opened. Within this different 128-gene, rank-4 family, the context Oracle improves by 2.25% for Tgfbr2 and 2.24% for Ifngr2

| Method | Overall | Random | Spatial block | Units |
|---|---|---|---|---|
| Uniform reference mean | 119.752 | 106.439 | 133.064 | 10 |
| Empirical reference | 42.941 | 21.716 | 64.166 | 10 |
| Learned anchor | 40.693 | 23.176 | **58.209** | 10 |
| **RADF** | **40.595** | **22.641** | 58.550 | 10 |

Table S1: Reference-source decomposition on the matched benchmark. Values are mean E-distance, and lower is better. The uniform reference mean repeats one centroid, whereas the learned anchor is population valued and balanced.

| Protocol | Comparison | Baseline ED | Candidate ED | Relative change | Units |
|---|---|---|---|---|---|
| Matched capacity | Reference decoder vs. relational flow | 35.666 | 35.915 | $+0.70\%$ | 4 |
| Prospective spatial | Anchor vs. RADF | 89.011 | 87.909 | $-1.24\%$ | 1 |
| Prospective random | Anchor vs. RADF | 15.811 | 15.351 | $-2.91\%$ | 1 |

Table S2: Capacity and prospective utility. Negative relative change denotes lower E-distance for RADF. The capacity and prospective rows use different protocols and are not pooled.

(Table S5). These achieved increments support context-dependent representational headroom; they are neither a theoretical ceiling nor leakage-safe model performance.

### RADF Remains Effective Across Increasing Spatial Shift

The external evaluations order conditions by increasing separation between the reported reference population and the evaluation context. E-distance values are comparable only within a dataset and preprocessing space. On leave-one-slide-out Perturb-map, RADF reduces E-distance by 54.2% relative to CONCERT, although scGen and direct empirical reuse are slightly lower in this particular track. On sealed same-slide SPAC-seq, RADF is the best learned method and is marginally better than empirical reuse. On the harder cross-replicate track, RADF remains better than CONCERT but trails scGen, indicating that replicate shift is not fully resolved by reference anchoring alone.

The cross-slide track contains five leave-one-slide-out Perturb-map units on 1,053 shared genes, each with 32 references from other slides. SPAC-seq uses 1,718 genes intersecting the frozen panel. Its sealed same-slide track contains 18 units with 32 references and 64 disjoint evaluation spots; the cross-replicate track contains nine units with 64 replicate-1 references and 128 replicate-2 evaluation spots. Radius 0.25 and checkpoint 300 are frozen from Perturb-map validation before either SPAC-seq track is scored. Table S6 reports the within-track E-distances for these three increasingly shifted settings; its rows are not placed on a shared numeric scale.

Spatially stratified diagnostics reveal an advantage that population-level E-distance alone cannot localize. Relative to CONCERT, RADF reduces SS-ED by 52.4% and contrast error by 32.8% on supported matched units. The reductions are 78.4% and 69.5%, respectively, on sealed same-slide SPAC-seq. On cross-replicate SPAC-seq, RADF lowers contrast error by 41.4% relative to CONCERT and also improves upon empirical reference reuse and scGen (Table S7).

### Biological and Regional Fidelity

Figure S1 first examines direct gene-level agreement on the ten matched task–split units. RADF has the strongest pair of macro summaries among the five methods, with Pearson $r = 0.971$ and MAE $= 0.102$, compared with $0.931/0.166$ for CONCERT, $0.966/0.104$ for scGen, $0.962/0.106$ for biolord, and $0.839/0.391$ for CellOT. Its points therefore remain most tightly aligned with the identity line in aggregate. This view tests recovery of held-out gene-wise means; it does not measure within-population variation, imply paired cells, or replace the primary distribution-level E-distance.

A complementary retrospective audit asks whether lower population error is accompanied by agreement with held-out biological response structure. It is not used for training, validation, or method selection. Expression is normalized to log-one-plus counts per 10,000, and each population response is the normalized target mean minus its matched KP/tumor source mean. Externally fixed mouse GO/MGI memberships from the frozen 2025-07-21 CausalInteractionBench snapshot define response to transforming growth factor beta (GO:0071559) for Tgfbr2 and response to type-II interferon (GO:0034341, GO:0060332, and GO:0071346) for Ifngr2. Intersecting these sets with each frozen gene panel leaves 23–28 genes per unit.

RADF attains the highest overall response Spearman (0.690), Top-50 recovery (0.638), and pathway-gene Spearman (0.841) among the evaluated methods (Table S8). The learned anchor has the lowest pathway-response error, with RADF close behind (0.146 versus 0.149), and both match the pathway-response direction in nine of ten units. This pattern is consistent with the intended division of labor: reference anchoring supplies most biological preservation, while bounded flow retains that structure during spatial refinement.

Regional pathway contrast is defined only for the three units satisfying the same spatial-support rule used by the coordinate audit. RADF improves on CONCERT and the learned anchor with mean absolute contrast error 0.199 and matches two of three directions (Table S9). The GO memberships are unsigned, so agreement measures fidelity to

| Unit | Candidate | Val. gain | Residual (%) | Saturation (%) | Eval. gain (%) | Status |
|---|---|---|---|---|---|---|
| 54-T-R | $\rho = .10, 300$ | 0.290 | 6.32 | 74.5 | 4.87 | nonzero |
| 54-T-S | $\rho = .25, 350$ | 2.223 | 6.09 | 52.0 | 0.39 | nonzero |
| 55-T-R | $\rho = .10, 300$ | 0.671 | 4.87 | 72.4 | -0.21 | nonzero |
| 55-T-S | anchor | 0.000 | 0.00 | – | 0.00 | fallback |
| 55-I-R | $\rho = .25, 400$ | 0.065 | 3.40 | 61.1 | 2.09 | nonzero |
| 55-I-S | $\rho = .10, 400$ | 0.956 | 2.24 | 62.2 | -0.38 | nonzero |
| 56-I-R | $\rho = .10, 350$ | 1.347 | 4.40 | 70.7 | 5.36 | nonzero |
| 56-I-S | $\rho = .25, 200$ | 3.474 | 4.41 | 46.0 | -1.47 | nonzero |
| 57-T-R | $\rho = .10, 200$ | 1.060 | 7.85 | 78.1 | 0.37 | nonzero |
| 57-T-S | $\rho = .25, 300$ | 2.025 | 6.35 | 40.6 | -0.73 | nonzero |

Table S3: Validation-only selection for the ten matched units. T/I denote Tgfbr2/Ifngr2 and R/S denote random/spatial-block splits. Validation gain is anchor E-distance minus selected-candidate E-distance. Evaluation gain is the percentage E-distance improvement over the anchor; it is reported only after selection is frozen. Saturation is the fraction of output entries on a tube boundary.

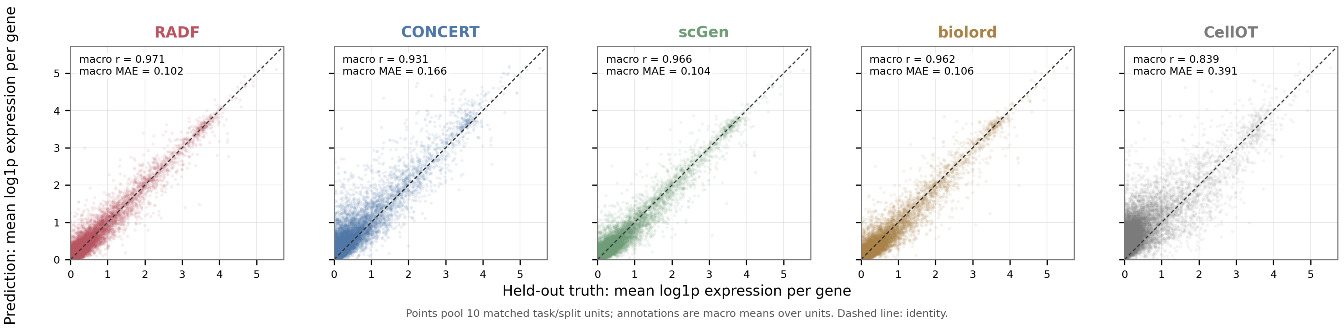


Figure S1: Gene-level agreement between held-out truth and frozen predictions on the ten matched Perturb-map task–split units. Each point is one gene in one unit and compares the mean of log-one-plus expression across the unpaired predicted and held-out populations, giving 20,000 points per method. Panels pool the points, whereas annotations are arithmetic macro means of the ten per-unit Pearson correlations and MAEs. The statistics are descriptive, do not enter model selection, and do not assess within-population variability. The dashed line denotes identity.

| Intervention | Pred. change (%) | Δ contrast error | Δ SS-ED |
|---|---|---|---|
| Permutation | 2.631 | +1.217 | -0.853 |
| Collapse | 2.338 | +1.132 | -0.639 |
| Region swap | 2.106 | +0.772 | -0.378 |

Table S4: Coordinate interventions over the three supported units. Positive contrast-error changes favor the original coordinates. SS-ED moves slightly in the opposite direction, showing that within-region distribution matching and regional separation capture different properties.

held-out response and does not imply pathway activation, inhibition, discovery, or perturbation-target causality. The target genes Tgfbr2 and Ifngr2 are absent from the frozen panels.

The independent sampling units are the five slide–perturbation task clusters, not the ten task–split rows or individual spots. Across 10,000 deterministic cluster-bootstrap draws, the paired RADF advantage over CONCERT is $[0.154, 0.376]$ for response Spearman and $[0.039, 0.479]$ for pathway response absolute error. These intervals are de-

| Episode | Global Oracle | Context Oracle | Gain |
|---|---|---|---|
| Tgfbr2 | 1.2918 | 1.2628 | 2.25% |
| Ifngr2 | 3.4175 | 3.3409 | 2.24% |

Table S5: Truth-assisted context headroom. Lower E-distance is better.

scriptive because only five clusters are available.

### Conservative Evaluation Prevents Unsupported Gains

RADF uses a deny-by-default evaluation boundary. Fitting and candidate generation read only query coordinates and reported reference expression. Candidate predictions are frozen before validation expression is opened; validation selects one candidate; evaluation expression is then read once for scoring. The verification results in Table S10 establish that this access order is reflected in the saved prediction paths.

For each new prediction path, evaluation expression is replaced by an extreme value and prediction is replayed. The ten-unit audit records maximum prediction change zero un-

| Evaluation | RADF | CONCERT | scGen | Empirical reference |
|---|---|---|---|---|
| Perturb-map cross-slide | 71.199 | 155.303 | **69.651** | 69.883 |
| SPAC-seq same-slide | **0.471** | 3.686 | 1.147 | 0.471 |
| SPAC-seq cross-replicate | 2.203 | 2.645 | **1.424** | 2.203 |

Table S6: External E-distance. Lower is better. Values must not be compared across rows because datasets and preprocessing spaces differ. Unrounded values determine bolding.

| Setting | Metric | CONCERT | RADF | Reduction |
|---|---|---|---|---|
| Matched supported | SS-ED | 138.404 | 65.910 | 52.4% |
| Matched supported | Contrast error | 83.300 | 55.998 | 32.8% |
| SPAC-seq same-slide | SS-ED | 3.910 | 0.844 | 78.4% |
| SPAC-seq same-slide | Contrast error | 0.652 | 0.199 | 69.5% |
| SPAC-seq cross-replicate | Contrast error | 0.511 | 0.300 | 41.4% |

Table S7: Spatially stratified diagnostics. Reduction is relative to CONCERT. The cross-replicate contrast errors for empirical reference and scGen are 0.376 and 0.377, respectively.

| Method | Response $\rho$ | Top-50 | Pathway error | Pathway-gene $\rho$ | Direction |
|---|---|---|---|---|---|
| CellOT | 0.332 | 0.306 | 0.227 | 0.576 | 9/10 |
| scGen | 0.625 | 0.546 | 0.251 | 0.784 | 9/10 |
| biolord | 0.586 | 0.450 | 0.315 | 0.742 | 8/10 |
| CONCERT | 0.423 | 0.330 | 0.406 | 0.569 | 5/10 |
| Learned anchor | 0.688 | 0.630 | **0.146** | 0.829 | 9/10 |
| **RADF** | **0.690** | **0.638** | 0.149 | **0.841** | 9/10 |

Table S8: Frozen matched-task biological fidelity over ten units. Response $\rho$ is Spearman correlation over the 2,000-gene perturbation-response vector. Top-50 is overlap with the largest absolute observed responses. Pathway error is absolute matched-set response-score error.

| Method | Contrast error | Direction |
|---|---|---|
| CellOT | 0.273 | 0/3 |
| scGen | 0.226 | 2/3 |
| biolord | **0.185** | 1/3 |
| CONCERT | 0.207 | 1/3 |
| Learned anchor | 0.201 | 2/3 |
| RADF | 0.199 | 2/3 |

Table S9: Regional pathway-response fidelity on the three supported units.

der this mutation, maximum replay error zero across 168 replays, finite nonnegative outputs, unchanged input hashes, and explicit retention of seven units that lack common spatial-stratum support. The prospective audits also verify balanced coupling marginals, anchor-tube containment, and dynamic relation recomputation. These checks support leakage isolation and structural validity; they do not turn descriptive small-sample intervals into confirmatory evidence.

### Supporting Definitions and Implementation Details

#### End-to-End RADF Procedure

Algorithm S1 makes RADF's three safeguards explicit. The reported population supplies both the balanced anchors and the flow-matching endpoint, projected integration bounds every nonzero refinement, and validation retains that refinement only when it improves upon the exact anchor. Candidate predictions for both validation and evaluation coordinates are frozen before validation expression is opened; held-out evaluation expression is not an algorithm input and is read only after selection is fixed.

#### Population and Spatial Metrics

For $U = \{u_i\}_{i=1}^N$ and $V = \{v_j\}_{j=1}^M$, the empirical V-statistic in aligned gene space is

$$D_{\mathrm{E}}(U,V) = \frac{2}{NM}\sum_{i=1}^{N}\sum_{j=1}^{M}\|u_i - v_j\|_2 - \frac{1}{N^2}\sum_{i=1}^{N}\sum_{i'=1}^{N}\|u_i - u_{i'}\|_2 - \frac{1}{M^2}\sum_{j=1}^{M}\sum_{j'=1}^{M}\|v_j - v_{j'}\|_2. \tag{S3}$$

Evaluation clamps only negative numerical round-off to zero. Lower values are better.

For spatial diagnostics, pooled query and truth coordinates are projected onto their first spatial principal axis and divided at the median. Let $\widehat{Y}_r$ and $Y_r$ denote prediction and truth in

| Audit | Split/source rederived | Mutation guard | Prediction replay | Metric recomputation |
|---|---|---|---|---|
| Matched capacity | yes | passed | passed | passed |
| Prospective spatial | yes | passed | passed | passed |
| Prospective random | yes | passed | passed | passed |
| Ten-unit spatial audit | coordinate-only strata | max change 0 | 168/168 | 119 rows |
| Biological fidelity | frozen source hashes | protocol before access | not applicable | 774 values |

Table S10: Compact verification receipt. The ten-unit audit independently recomputed E-distance, PCC, MAE, SS-ED, and spatial-contrast error. All listed audits have verified or passed status.

stratum $r$. Then

$$\text{SS-ED} = \sum_{r\in\{0,1\}} \frac{|Y_r|}{|Y_0|+|Y_1|} D_{\mathrm{E}}(\widehat{Y}_r, Y_r). \tag{S4}$$

Spatial-contrast error is the absolute difference between predicted and observed cross-stratum separation:

$$\left| D_{\mathrm{E}}(\widehat{Y}_0, \widehat{Y}_1) - D_{\mathrm{E}}(Y_0, Y_1) \right|. \tag{S5}$$

Each of the four strata must contain at least four samples.

### Log-Domain Balanced Allocation

Writing $\alpha$ and $\beta$ for row and column log-scalings, one Sinkhorn iteration applies

$$\alpha_i \leftarrow -\log n - \mathrm{LSE}_j(L_{\phi,ij} + \beta_j), \tag{S6}$$

$$\beta_j \leftarrow -\log m - \mathrm{LSE}_i(L_{\phi,ij} + \alpha_i), \tag{S7}$$

$$W_{\phi,ij} \leftarrow n \exp(L_{\phi,ij} + \alpha_i + \beta_j). \tag{S8}$$

Here LSE is log-sum-exp. With $\overline{W}_{\phi,ij} = \max(W_{\phi,ij}, 10^{-12})$, the allocation regularizer is

$$H(W_\phi) = -\frac{1}{n}\sum_{i=1}^{n}\sum_{j=1}^{m} \overline{W}_{\phi,ij} \log \overline{W}_{\phi,ij}. \tag{S9}$$

### Endpoint Normalization and Projected Integration

Let $\mu = \mathrm{mean}(\mathrm{concat}(A_p, R_p))$ and $\sigma = \max(\mathrm{std}(\mathrm{concat}(A_p, R_p)), 1)$. The standardized coupling inputs are

$$\overline{A} = (A_p - \mu)/\sigma, \qquad \overline{R} = (R_p - \mu)/\sigma. \tag{S10}$$

The detached normalization scales used by the losses are

$$s_v = \max\left(\sqrt{\mathrm{mean}((B_p - A_p)^2)}, 1\right). \tag{S11}$$

$$s_E = \max(D_{\mathrm{E}}(A_p, R_p), 1). \tag{S12}$$

For anchor $A_p$ and radius $\rho$, the fixed bounds are

$$\mathrm{lower}_\rho = \max(0, A_p - \rho(A_p + \mathbf{1})). \tag{S13}$$

$$\mathrm{upper}_\rho = A_p + \rho(A_p + \mathbf{1}). \tag{S14}$$

All maxima, divisions, and clipping operations involving matrices are element-wise.

| Component | Setting |
|---|---|
| Anchor optimization | 400 Adam steps, lr 0.05 |
| Anchor Sinkhorn | 160 iterations |
| Entropy coefficient | $\lambda_H = 0.05$ |
| Hidden dimension | $d = 32$ |
| Training coupling | $\epsilon = 0.5$, 120 iterations |
| Flow optimization | 400 Adam steps, lr $10^{-3}$ |
| Reference-loss weight | 0.5 |
| Gradient clipping | norm 5 |
| Euler integration | 8 steps |
| Candidate radii | $\{0.10, 0.25\}$ |
| Checkpoints | $\{200, 300, 350, 400\}$ |

Table S11: Frozen RADF hyperparameters for the matched benchmark.

### Frozen Hyper-parameters

Table S11 consolidates the frozen settings used by Algorithm S1. The two-radius, four-checkpoint grid is evaluated only through validation E-distance, and none of these settings is chosen using held-out evaluation expression.

The anchor, coupling, and loss scales are detached from gradient updates. Separate models are trained for the two nonzero radii because projection changes the reference loss. The exact anchor is included as candidate $c_0$. Ties among improving nonzero candidates prefer the smaller radius and then the earlier checkpoint. PCC and MAE are descriptive and do not enter fitting, selection, or fallback.

**Algorithm S1** RADF with reference anchoring, bounded refinement, and exact fallback

**Require:** Reported population $R_p$; query coordinates $S^{\mathrm{val}}, S^{\mathrm{eval}}$
  **Fixed:** $\mathcal{R} = \{0.10, 0.25\}$, $\mathcal{K} = \{200, 300, 350, 400\}$, $K_{\mathrm{int}} = 8$
**Ensure:** Frozen evaluation prediction $\widehat{Y}_p^{\mathrm{eval}}$
**for** $q \in \{\mathrm{val}, \mathrm{eval}\}$ **do**
  $n_q \leftarrow |S^q|$
  $W_\phi^q \leftarrow \textsc{FitBalancedAnchor}(R_p, n_q)$
  $A_p^q \leftarrow W_\phi^q R_p$
**end for**
$\Omega \leftarrow \{c_0\}$
$\widehat{Y}_{c_0}^q \leftarrow A_p^q$ for $q \in \{\mathrm{val}, \mathrm{eval}\}$
$\Gamma_p \leftarrow \textsc{FixedSinkhorn}(A_p^{\mathrm{eval}}, R_p)$
$B_p^{\mathrm{eval}} \leftarrow \Gamma_p R_p$; compute and detach $s_v, s_E$
**for** $\rho \in \mathcal{R}$ **do**
  Initialize $\theta_\rho$ identically with zero output weights
  **for** $k = 1, \ldots, 400$ **do**
    Sample $t \sim \mathrm{Uniform}([0,1]^{n_{\mathrm{eval}}})$
    $Z(t) \leftarrow (1-t) \odot A_p^{\mathrm{eval}} + t \odot B_p^{\mathrm{eval}}$
    Recompute state-dependent relations from $(Z(t), S^{\mathrm{eval}})$
    Evaluate $\mathcal{L}_{\mathrm{CFM}}$ on $Z(t)$
    $\widetilde{Y} \leftarrow F_{\theta_\rho,\rho}(S^{\mathrm{eval}}, R_p; A_p^{\mathrm{eval}})$
    $\mathcal{L}_{\mathrm{ref}} \leftarrow D_{\mathrm{E}}(\widetilde{Y}, R_p)/s_E$
    Update $\theta_\rho$ using $\mathcal{L}_{\mathrm{CFM}} + 0.5\mathcal{L}_{\mathrm{ref}}$
    Clip the gradient norm to 5
    **if** $k \in \mathcal{K}$ **then**
      Freeze $\theta_{\rho,k}$; set $c = (\rho, k)$
      $\Omega \leftarrow \Omega \cup \{c\}$
      **for** $q \in \{\mathrm{val}, \mathrm{eval}\}$ **do**
        $\widehat{Y}_c^q \leftarrow F_{\theta_{\rho,k},\rho}(S^q, R_p; A_p^q)$
      **end for**
    **end if**
  **end for**
**end for**
Freeze $\Omega$; only now open $Y_p^{\mathrm{val}}$
**for** $c \in \Omega$ **do**
  $e_c \leftarrow D_{\mathrm{E}}(\widehat{Y}_c^{\mathrm{val}}, Y_p^{\mathrm{val}})$
**end for**
$c^+ \leftarrow \arg\min_{c \in \Omega \setminus \{c_0\}} e_c$
  ▷ Break ties by smaller $\rho$, then earlier $k$
**if** $e_{c^+} < e_{c_0}$ **then**
  $c^\star \leftarrow c^+$
**else**
  $c^\star \leftarrow c_0$ ▷ exact anchor fallback
**end if**
$\widehat{Y}_p^{\mathrm{eval}} \leftarrow \widehat{Y}_{c^\star}^{\mathrm{eval}}$; freeze it
Open $Y_p^{\mathrm{eval}}$ once and score the frozen prediction
